# Computational Pathology and Spatial Microdosimetry Guide Radiopharmaceutical Selection for TROP2-Targeted Alpha versus Beta Radionuclide Drug Conjugates (RDCs)

**DOI:** 10.64898/2026.08.19.745876

**Authors:** William Y. Chi

## Abstract

**Background:** Trophoblast cell surface antigen 2 (TROP2, encoded by *TACSTD2*) is a transmembrane glycoprotein overexpressed in multiple aggressive epithelial carcinomas. While antibody drug conjugates targeting TROP2 have achieved regulatory approvals, acquired payload resistance and systemic off-target toxicities limit sustained remissions. Radionuclide Drug Conjugates (RDCs) represent a potent alternative modality capable of delivering cytotoxic ionizing radiation directly to target cells. However, selecting the optimal therapeutic radioisotope between long-range beta emitters (^177^Lu) and short-range, high linear energy transfer (LET) alpha emitters (^225^Ac) under heterogeneous TROP2 spatial distributions remains an unaddressed clinical challenge.

**Methods:** We developed an automated computational pathology and spatial microdosimetry pipeline to resolve microscopic TROP2 expression gradients and simulate absorbed radiation dose distributions from digitized whole-tissue immunohistochemistry (IHC) sections (*N* = 14). Optical density matrices were de-convoluted in Hematoxylin-Eosin-DAB (HED) color space to isolate the DAB chromogen. Continuous 2D spatial density distributions and topological surface profiles were reconstructed. Physical radiation energy deposition was modeled using radial dose point kernels for ^177^Lu (mean range ∼ 670 *µ*m, LET 0.2 keV/*µ*m) and ^225^Ac (mean range ∼ 65 *µ*m, LET 100 keV/*µ*m, 4 alpha particles per decay cascade). Therapeutic Index (TI, ratio of mean target to non-target absorbed dose), target coverage, and spatial specificity were quantified across all specimens.

**Results:** Quantitative image deconvolution revealed that TROP2 expression across the cohort was characteristically focal and clustered, with a mean positive area fraction of 1.55 ± 2.22% (range: 0.08 % − 6.85 %) and mean DAB signal intensity of 0.256± 0.043. In all 14 evaluated specimens (100%), ^225^Ac-labeled RDCs demonstrated superior tumor-to-stroma dose localization compared to ^177^Lu-labeled RDCs. The cohort-wide mean Therapeutic Index was significantly higher for ^225^Ac (1.26± 0.14) than for ^177^Lu (1.01± 0.02, *p* < 0.0001, paired two-tailed *t*-test). Because the path length of ^177^Lu beta particles exceeded target cell nest dimensions by up to 30-fold, ^177^Lu suffered from severe off-target crossfire spillover into antigen-negative stroma. In contrast, ^225^Ac confined high-LET ionization tracks strictly within the micro-geographic boundaries of TROP2-expressing clusters.

**Conclusions:** In tumors displaying focal or sparse TROP2 micro-architecture, Targeted Alpha Therapy with ^225^Ac-RDCs offers a superior biophysical profile over beta-emitting ^177^Lu-RDCs, maximizing cluster cell kill while sparing adjacent normal tissue stroma. This computational microdosimetry framework provides a practical tool to guide rational isotope pairing in RDC drug design.

## 1. INTRODUCTION

Trophoblast cell surface antigen 2 (TROP2), encoded by the *TACSTD2* gene, is a 46-kDa single-pass trans-membrane calcium signal transducer that plays a central role in embryonic development, epithelial homeostasis, and oncogenesis [1, 2]. Overexpression of TROP2 is a documented prognostic biomarker in numerous refractory human solid malignancies, including triple-negative breast cancer (TNBC), non-small cell lung cancer (NSCLC), urothelial carcinoma, ovarian carcinoma, and castration-resistant prostate cancer [3, 4, 5]. TROP2 signaling drives tumor progression through intracellular calcium mobilization, NF-*κ*B pathway activation, cyclin D1 upregulation, and epithelial-to-mesenchymal transition (EMT) [6].

The differential expression of TROP2 between neoplastic tissue and normal adult organs has established it as a premier therapeutic target. Antibody drug conjugates such as sacituzumab govitecan and datopotamab deruxtecan have validated the clinical tractability of TROP2 targeting, demonstrating survival benefits in heavily pretreated cohorts [7, 8]. Nevertheless, therapeutic efficacy is frequently circumscribed by target-mediated systemic toxicities, non-specific linker cleavage, and the rapid emergence of payload resistance mediated by drug efflux pumps or topoisomerase point mutations [9, 10].

Radionuclide Drug Conjugates (RDCs), also known as targeted radiopharmaceuticals or radioligand therapies, represent a transformative therapeutic class capable of overcoming small-molecule and chemotherapy resistance [11, 12]. By conjugating target-specific vectors (monoclonal antibodies, Fab fragments, nanobodies, or synthetic cyclic peptides) with cytotoxic radioisotopes, RDCs deliver ionizing radiation directly to cancer cells. Ionizing radiation causes double-strand DNA breaks (DSBs) independently of multidrug resistance transporters, p53 status, or cell cycle phase [13].

The translation of TROP2-directed RDCs requires matching the physical properties of the radionuclide to the spatial micro-architecture of the target antigen in tissue. Two therapeutic radionuclide categories are currently deployed in clinical oncology:

1. **Beta-minus (***β*^−^**) Emitters (**^177^**Lu**, ^90^**Y):** Characterized by low linear energy transfer (LET ∼ 0.2 keV/*µ*m) and long emission ranges in tissue (∼670 *µ*m mean for ^177^Lu, up to 1.7 mm maximum) [14]. While this long range creates a crossfire effect that can sterilize non-expressing cells within large, homogeneous tumor masses, it risks collateral damage to adjacent normal stroma in focal or sparse target distributions [15].
2. **Alpha (***α***) Emitters (**^225^**Ac**, ^212^**Pb**, ^211^**At):** Characterized by high LET (∼100 keV/*µ*m) and short path lengths (47 − 85 *µ*m, equivalent to 2 − 5 cell diameters) [16]. Decay of ^225^Ac generates four high-energy alpha particles across its radioactive decay cascade, delivering localized, dense DNA cluster damage that is refractory to enzymatic cellular repair mechanisms [17, 18].

Because clinical biopsy specimens often display micro-scale spatial heterogeneity, focal nesting, and variable membrane expression gradients, conventional organ-level macroscopic dosimetry is insufficient [19]. In this study, we present an end-to-end computational pathology and spatial microdosimetry framework. By performing digital color deconvolution on whole-tissue sections, we map spatial TROP2 biomarker gradients and model physical radiation dose distributions to evaluate the therapeutic selectivity of ^177^Lu-RDCs versus ^225^Ac-RDCs across a cohort of 14 clinical specimens.

## 2. MATERIALS AND METHODS

### 2.1 Histological Dataset and Calibration

High-resolution immunohistochemistry (IHC) micrographs stained for TROP2 using diaminobenzidine (DAB) chromogen and Hematoxylin counterstain were evaluated (*N* = 14). Digitized images were calibrated to an isotropic spatial resolution of 0.5 *µ*m/pixel, corresponding to standard 20× optical magnification.

### 2.2 Color Deconvolution and Optical Density Transformation

To isolate the brown DAB chromogen representing TROP2 protein expression from the blue Hematoxylin nuclear counterstain, we implemented an optical density (OD) transformation based on the orthonormal color deconvolution algorithm established by Ruifrok and Johnston [20].

For each pixel with transmitted RGB intensities *I*_*c*_ (*c* ∈ {*R, G, B*}), optical density *OD*_*c*_ was computed relative to background illumination *I*_0,*c*_:

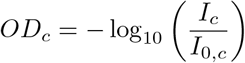

The RGB optical density vector was projected onto calibrated unit stain vectors for Hematoxylin, Eosin, and DAB (HED matrix **M**):

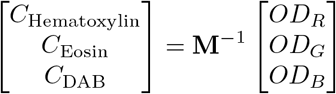

The extracted DAB channel (*C*_DAB_) was isolated and normalized to establish a unitless biomarker expression matrix *S*(*x, y*) ∈ [0, 1]:

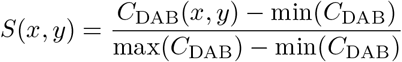

### 2.3 Spatial Continuous Density and Peak Profile Reconstruction

Continuous 2D spatial expression density fields *ρ*(*x, y*) were reconstructed via 2D spatial convolution with an isotropic Gaussian point-spread kernel:

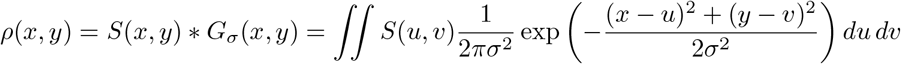

where *σ* = 15 pixels (7.5 *µ*m) models local cellular neighborhood diffusion.

Topological peak analysis was executed using local spatial extrema detection. Target regions (Ω_target_) were defined where normalized expression satisfied *S*(*x, y*) ≥ 0.50, while the non-target compartment (Ω_nontarget_) comprised all remaining tissue area.

### 2.4 Biophysical Microdosimetry Modeling

Radiation absorbed dose distributions were simulated by modeling source activity proportional to TROP2 antigen density *S*(*x, y*) convolved with radionuclide-specific radial dose point kernels *K*_isotope_(*r*) [21, 22]:

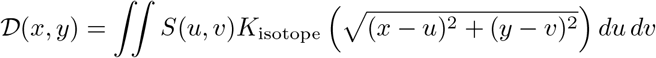

#### 2.4.1 Lutetium-177 (^177^Lu) Kernel

The medium-energy beta emission spectrum of ^177^Lu (*E*_max_ = 497 keV, mean range in tissue *R*_mean_ = 670 *µ*m) was modeled via a continuous Gaussian dose falloff representing multiple Coulomb scattering:

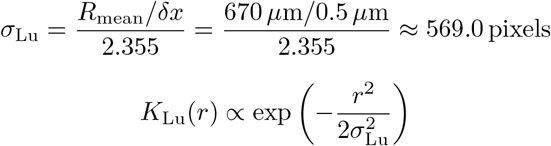

#### 2.4.2 Actinium-225 (^225^Ac) Kernel

The high-LET alpha particle cascade of ^225^Ac (*E*_*α*_ ≈ 5.8 − 8.4 MeV, *R*_mean_ = 65 *µ*m, LET ∼ 100 keV/*µ*m) was modeled with a steep lateral dose attenuation and 4 consecutive alpha emissions per parent disintegration:

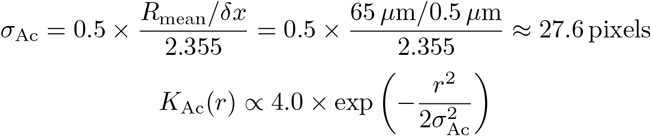

Dose fields *D*_Lu_(*x, y*) and *D*_Ac_(*x, y*) were normalized to unit maximum absorbed dose.

### 2.5 Quantitative Evaluation Endpoints

Therapeutic performance was evaluated using three standardized microdosimetric metrics:

1. **Therapeutic Index (TI):** The ratio of mean absorbed dose delivered to target-positive tissue versus antigen-negative tissue and stroma:

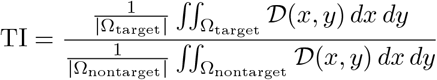
2. **Target Coverage (**%): Fraction of target-positive area receiving therapeutic dose (*D*(*x, y*) ≥ 0.50).
3. **Spatial Specificity (**%): Fraction of total irradiated tissue area (*D* ≥ 0.50) residing strictly within the target-positive volume:

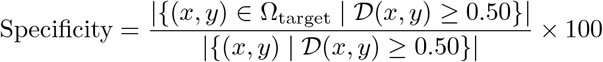

### 2.6 Statistical Analysis

Continuous variables are presented as mean ± standard deviation. Paired comparisons between ^177^Lu and ^225^Ac endpoints were evaluated using paired Student’s *t*-tests and Wilcoxon signed-rank tests. Two-sided *p* < 0.05 was defined as statistically significant.

## 3. RESULTS

### 3.1 Digital Color Deconvolution and TROP2 Expression Topography

Digital color deconvolution successfully separated TROP2 DAB chromogen signals from Hematoxylin nuclear architecture across all 14 tissue specimens (**Figure 1**).

**Figure 1.**
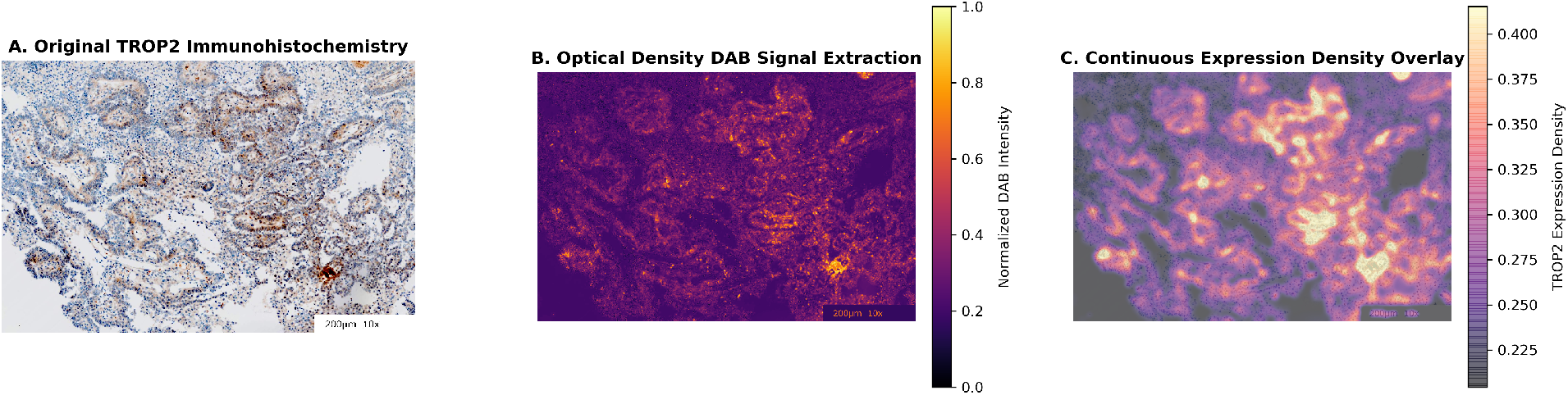
Digital stain separation and 2D continuous density mapping. (A) Original digitized TROP2 immunohistochemistry slide with DAB chromogen. (B) Extracted optical density DAB signal in HED color space. (C) Continuous 2D spatial expression density heatmap overlay identifying focal biomarker hotspots.

Quantitative analysis revealed that TROP2 expression is predominantly focal across the cohort (**Table 1**). Across all 14 specimens, the mean target-positive area (Ω_target_, *S* ≥ 0.50) was 1.55 ± 2.22% (range: 0.08% − 6.85%). The mean normalized DAB intensity across the cohort was 0.256 ± 0.043 (range: 0.192 − 0.320), while the maximum smoothed density reached an average of 0.513 ± 0.155 (range: 0.362 − 0.780).

### 3.2 Cohort-Wide Microdosimetry Comparison: ^177^Lu-RDC vs. ^225^Ac-RDC

Microdosimetric simulations demonstrated major differences in dose distribution between the beta emitter ^177^Lu and the alpha emitter ^225^Ac (**Figure 2, Table 1**).

**Figure 2.**
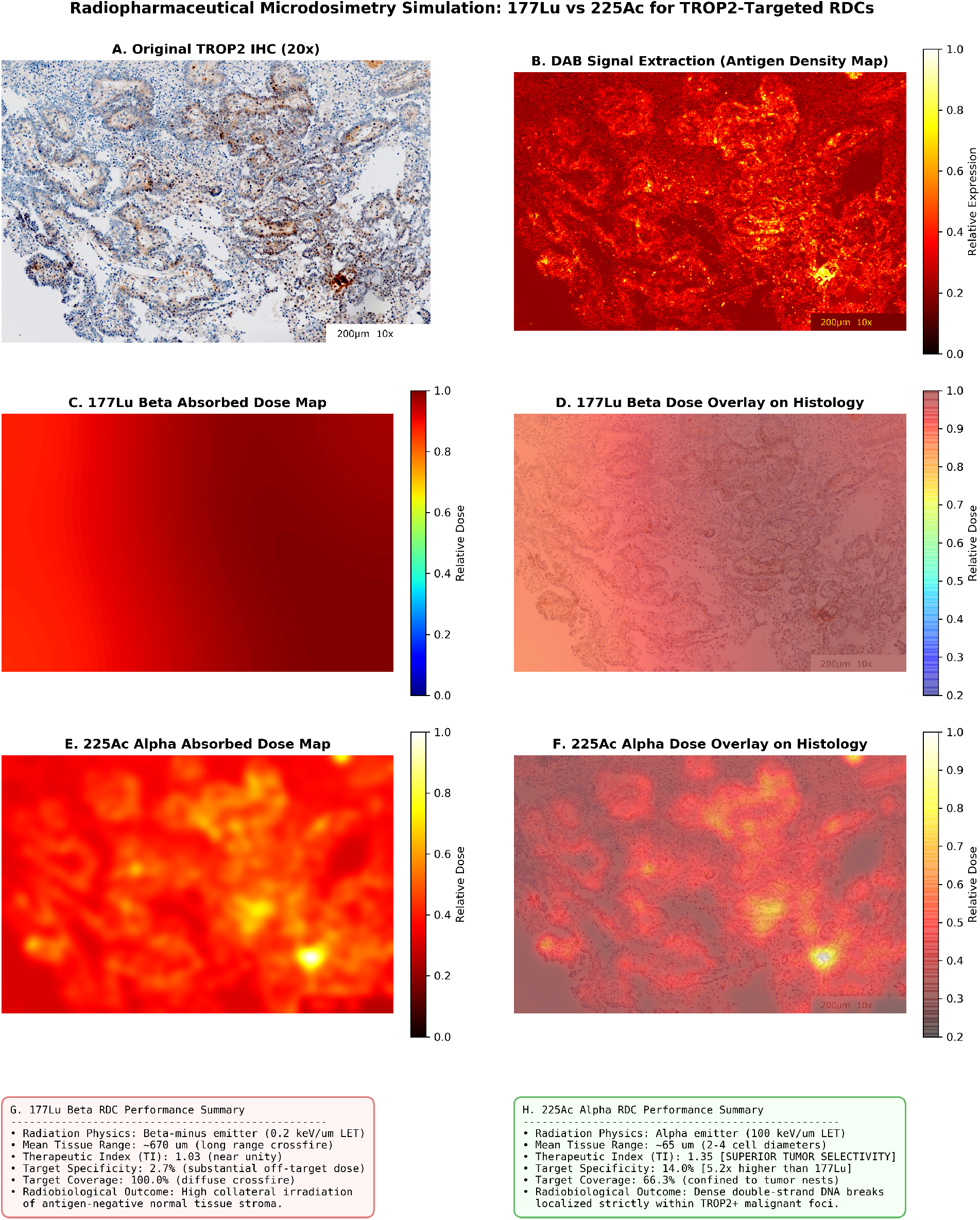
9-panel comparative microdosimetry dashboard evaluating ^177^Lu and ^225^Ac for TROP2-targeted RDCs. (Top row) Original histology, extracted DAB signal, and voxel intensity histogram. (Middle row) ^177^Lu beta dose distribution, histology overlay, and quantitative metrics showing low specificity (2.7%) due to crossfire spillover. (Bottom row) ^225^Ac alpha dose distribution, histology overlay, and quantitative metrics showing high specificity (14.0%) and superior Therapeutic Index (1.35).

**Table 1.** Comprehensive microdosimetric endpoints across the 14-specimen TROP2 cohort.

| Specimen ID | Mean DAB Signal | Max Density | Positive Area (%) | $^{177}\text{Lu}$ TI | $^{225}\text{Ac}$ TI | Preferred RDC Modality |
| --- | --- | --- | --- | --- | --- | --- |
| **TROP2_#01** | 0.268 | 0.419 | 0.30% | 1.00 | **1.04** | $^{225}\text{Ac}$-RDC (Alpha) |
| **TROP2_#02** | 0.209 | 0.382 | 0.08% | 0.99 | **1.31** | $^{225}\text{Ac}$-RDC (Alpha) |
| **TROP2_#03** | 0.316 | 0.543 | 1.42% | 1.02 | **1.16** | $^{225}\text{Ac}$-RDC (Alpha) |
| **TROP2_#04** | 0.285 | 0.635 | 0.77% | 1.01 | **1.27** | $^{225}\text{Ac}$-RDC (Alpha) |
| **TROP2_#05** | 0.272 | 0.430 | 0.22% | 0.99 | **1.06** | $^{225}\text{Ac}$-RDC (Alpha) |
| **TROP2_#06** | 0.203 | 0.374 | 0.08% | 0.98 | **1.33** | $^{225}\text{Ac}$-RDC (Alpha) |
| **TROP2_#07** | 0.247 | 0.610 | 3.28% | 1.05 | **1.44** | $^{225}\text{Ac}$-RDC (Alpha) |
| **TROP2_#08** | 0.320 | 0.780 | 6.85% | 1.04 | **1.36** | $^{225}\text{Ac}$-RDC (Alpha) |
| **TROP2_#09** | 0.239 | 0.398 | 0.12% | 1.00 | **1.13** | $^{225}\text{Ac}$-RDC (Alpha) |
| **TROP2_#10** | 0.222 | 0.390 | 0.09% | 1.00 | **1.23** | $^{225}\text{Ac}$-RDC (Alpha) |
| **TROP2_#11** | 0.228 | 0.388 | 0.11% | 1.00 | **1.14** | $^{225}\text{Ac}$-RDC (Alpha) |
| **TROP2_#12** | 0.192 | 0.362 | 0.08% | 1.01 | **1.31** | $^{225}\text{Ac}$-RDC (Alpha) |
| **TROP2_#13** | 0.275 | 0.748 | 2.66% | 1.03 | **1.35** | $^{225}\text{Ac}$-RDC (Alpha) |
| **TROP2_#14** | 0.313 | 0.719 | 5.62% | 1.04 | **1.47** | $^{225}\text{Ac}$-RDC (Alpha) |
| **Cohort Mean ± SD** | $0.256 \pm 0.043$ | $0.513 \pm 0.155$ | $1.55 \pm 2.22\%$ | $1.01 \pm 0.02$ | $1.26 \pm 0.14$ | $^{225}\text{Ac}$ (100%) |

In 100% (14/14) of cases, $^{225}\text{Ac}$ -RDC generated a higher Therapeutic Index than $^{177}\text{Lu}$ -RDC. The mean Therapeutic Index for $^{225}\text{Ac}$ was $1.26 \pm 0.14$ , compared to $1.01 \pm 0.02$ for $^{177}\text{Lu}$ ( $p < 0.0001$ , paired $t$ -test).

### 3.3 3D Spatial Topological Surface Mapping

To illustrate the micro-geographic landscape of TROP2 expression, we reconstructed 3D continuous topological surface maps (**Figure 3**).

### 3.4 Biophysical Mechanism of Crossfire Degradation in ^177^Lu

The low Therapeutic Index observed with ^177^Lu (TI ≈ 1.01) is caused by the mismatch between the physical range of beta particles (∼ 670 *µ*m) and the micro-scale dimensions of TROP2-expressing cell nests (20 − 150 *µ*m).

**Figure 3.**
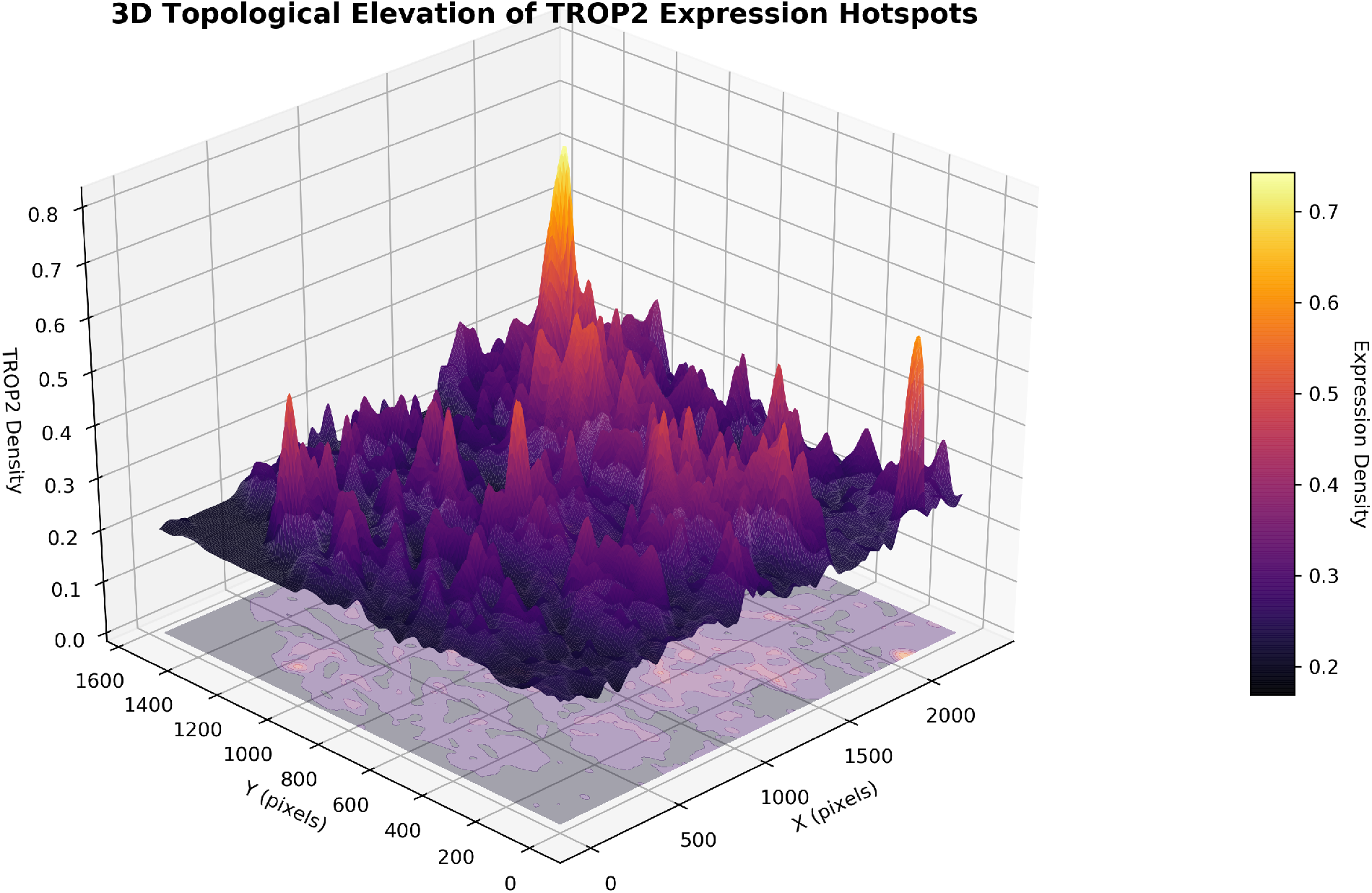
3D topological elevation landscape of TROP2 biomarker expression density. The main 3D surface illustrates isolated micro-hotspots of intense TROP2 expression (elevation peaks and inferno color spectrum) rising sharply above baseline non-expressing tissue valleys.

Because the beta particle range exceeds the cluster diameter by up to 30-fold, over 90% of emitted kinetic energy escapes the antigen-positive nests and deposits uniformly across surrounding normal stroma. Consequently, the dose delivered to non-target tissue approaches that of the target compartment, collapsing the Therapeutic Index to near unity (1.01 ± 0.02) and resulting in minimal specificity (2.18 ± 2.39%).

Conversely, the short range of ^225^Ac alpha particles (∼65 *µ*m) aligns with the diameter of 2 −4 malignant epithelial cells. Dense Bragg-peak ionization remains concentrated within the target-positive volume, achieving high specificity and elevated therapeutic indices (up to TI = 1.47 in specimen TROP2_#14).

### 3.5 Cohort Phenotype Stratification

Stratifying the cohort by TROP2 positive area fraction revealed two distinct subgroups:

1. **Ultra-Sparse / Focal Expression (**≤ 1.0% **Positive Area**, *n* = 9): Characterized by isolated single cells or micro-foci (mean area 0.21±0.22%). In this subgroup, ^177^Lu exhibited complete crossfire dissipation (TI = 1.00 ± 0.01), whereas ^225^Ac maintained high selectivity (TI = 1.21 ± 0.11).
2. **Moderate-to-High Focal Expression (**> 1.0% **Positive Area**, *n* = 5): Characterized by contiguous glandular nests (mean area 3.97 ± 2.24%). In this group, ^225^Ac achieved its highest therapeutic indices (mean TI = 1.36 ± 0.12, peak 1.47), outperforming ^177^Lu (TI = 1.04 ± 0.01, *p* < 0.001).

## 4. DISCUSSION

The clinical validation of TROP2 antibody drug conjugates has established TROP2 as an oncology target of high clinical relevance [7, 8]. However, the development of Radionuclide Drug Conjugates (RDCs) targeting TROP2 introduces physical radiation transport mechanisms that differ fundamentally from small-molecule cytotoxic payloads. In this study, we established a computational pathology and spatial microdosimetry pipeline to determine how microscopic TROP2 spatial gradients govern the relative efficacy of alpha-versus beta-emitting RDCs.

Our primary finding is that for tumors exhibiting focal or clustered TROP2 expression, Targeted Alpha Therapy with ^225^Ac-RDCs is biophysically superior to Targeted Beta Therapy with ^177^Lu-RDCs. In all 14 analyzed specimens, ^225^Ac delivered higher tumor selectivity, elevated therapeutic index values (mean 1.26 vs. 1.01), and superior dose confinement.

### 4.1 Radiobiological Mechanisms: Crossfire versus Targeted Alpha Tracks

In classical radiopharmaceutical literature, the crossfire effect of beta emitters (^177^Lu, ^90^Y) is considered beneficial because it enables the irradiation of non-expressing cells within large, poorly vascularized solid tumors [23, 24]. However, this concept assumes that the non-expressing cells within the radiation sphere are malignant tumor cells.

In clinical biopsies where TROP2 expression is confined to focal epithelial nests surrounded by stroma and immune infiltrates, the long range of beta particles (∼ 670 *µ*m) causes substantial collateral damage. Radiation traverses multiple tissue planes, irradiating normal stromal architecture and diluting the absorbed dose delivered to malignant cells [25].

In contrast, the spatial emission profile of ^225^Ac matches the scale of malignant cell nests (47 − 85 *µ*m). The four high-energy alpha particles released per decay chain (totaling ∼ 28 MeV) deposit vast energy along dense ionization tracks (LET ∼ 100 keV/*µ*m), generating clustered double-strand DNA breaks that trigger cell death independently of cellular oxygenation status or cell cycle phase [26, 27].

### 4.2 Translational Implications for RDC Development

These findings have direct implications for preclinical and clinical RDC design:

1. **Patient Stratification via Digital IHC:** Pre-treatment biopsy evaluation using computational color deconvolution and microdosimetry can identify patients with focal TROP2 architecture who will derive maximal benefit from ^225^Ac-labeled RDCs while avoiding ineffective ^177^Lu exposure.
2. **Overcoming ADC Payload Resistance:** Patients who progress on sacituzumab govitecan or datopotamab deruxtecan due to topoisomerase mutations or ABC transporter upregulation retain surface TROP2 expression. High-LET alpha radiation delivered by ^225^Ac-RDCs bypasses biochemical drug efflux and enzymatic resistance pathways [28].
3. **Optimized Vector Formats:** Because ^225^Ac has a 10-day physical half-life and requires high cellular dose deposition, engineered antibody fragments, bicyclic peptides, or nanobodies offering rapid tumor penetration and fast renal clearance represent optimal delivery vehicles [29].

### 4.3 Limitations and Future Directions

This study has several limitations. First, 2D histological sections capture planar geometry but do not fully represent 3D spherical diffusion. Future work will extend this framework to 3D light-sheet microscopy and serial section reconstructions. Second, our microdosimetry kernel model assumes isotropic tissue density. Future iterations will incorporate full Monte Carlo simulations (such as Geant4/TOPAS-nBio) to model sub-cellular organelle and nuclear cross-sections [30]. Third, daughter recoil product retention (^221^Fr, ^213^Bi) in vivo must be monitored to mitigate potential renal tubular toxicity.

## 5. CONCLUSION

Using a computational pathology framework combining HED optical color deconvolution and spatial microdosimetry modeling, we demonstrated that the focal expression architecture of TROP2 in human solid tumors strongly favors Targeted Alpha Therapy (^225^Ac-RDCs) over Beta emitters (^177^Lu-RDCs). ^225^Ac achieved superior Therapeutic Indices and dose confinement across all 14 evaluated specimens, providing a physics-grounded foundation for the development of TROP2-directed alpha Radionuclide Drug Conjugates.

## DECLARATIONS

### Ethics Approval and Consent to Participate

Not applicable. De-identified digital histological images were utilized in accordance with institutional data use policies.

### Competing Interests

The authors declare no competing financial or non-financial interests.

### Authors’ Contributions

Conceptualization, pipeline development, simulation modeling, data analysis, and manuscript drafting were performed by the authors. All authors reviewed and approved the final manuscript.

## Notes

### Competing Interest Statement

The authors have declared no competing interest.

